# Trial-by-trial fluctuations in post-stimulus attention during memory encoding predict subsequent associative context memory performance

**DOI:** 10.1101/2021.02.16.431454

**Authors:** Abdelhalim Elshiekh, M. Natasha Rajah

## Abstract

Episodic memory formation rate varies over time partly due to fluctuations in attentional state during memory encoding. Emerging evidence suggests that fluctuations in pre- and/or post-stimulus attention during encoding impact subsequent memory performance. It remains unclear how these fluctuations may differentially impact the subsequent retrieval of items alone, compared to items + their contextual details (associative context memory). In this study, we explored this in 30 healthy younger adults (21-34 years old). We developed the *Montreal Attention at Encoding* (MAET) task where on each encoding trial, participants responded as quickly as possible to a central fixation cross that expanded in size after a random duration. They then had to encode a picture of an object and its spatial location. Memory for the object-location associations was tested during retrieval. Response time (RT) to the fixation cross presented prior to each object gauged pre-stimulus attention levels on a trial-by-trial basis, while RT to the fixation cross that ensued each object indexed post-stimulus attention levels. Within-subject logistic regressions were used to predict context and item memory performance from pre- and post-stimulus RTs. Results revealed that encoding pre-stimulus attentional levels did not differentially predict context vs. item memory. However, post-stimulus RTs did predict subsequent context retrieval such that, longer post-stimulus RT to the fixation was related to poorer subsequent context retrieval. This study introduces a novel paradigm for investigating the impact of attentional state at encoding on subsequent memory performance and indicate a link between post-stimulus delays in attention-related RT and associative encoding success.

## 1. Introduction

Learning and memory hinge on the interaction of various underlying operations and processes. For instance, while intuition indicates that it is necessary to focus our attention during a lecture or in a classroom in order to improve encoding and subsequent recall of information, our attention constantly fluctuates between ‘good’ and ‘bad’ states (Esterman et al., 2013, 2014; Rosenberg et al., 2013). These moment-to-moment fluctuations in attentional state during encoding may therefore impact episodic memory performance, or our ability to recall an event and its spatiotemporal context (Markant et al., 2014; Tulving, 2002). Indeed, for any given individual, the rate of learning and memory formation varies over time in part due to fluctuations in the neural signals underpinning an individual’s current level of attention (Fernandez et al., 1999; Yoo et al., 2012). In the present study, we propose a novel paradigm to investigate whether fluctuations in attentional state preceding and following an episodic event to be encoded, predicts whether it would be recalled along with its contextual details.

Previous studies investigating the role of attention on episodic memory encoding have mostly relied on dual-task or divided attention paradigms (Anderson et al., 1998; Craik et al., 1996; 2018; Naveh-Benjamin et al., 1998; Troyer et al., 1999; Troyer & Craik, 2000). These paradigms investigate the stages of memory processing most prone to disruption in the face of distraction, and therefore those that are most in need of attentional resources under normal conditions (Hannula, 2018). In divided attention paradigms, participants are asked to encode a list of words while simultaneously engaging in a secondary task (e.g., monitoring certain tones) (Anderson et al., 1998; Kensinger et al., 2003; Park et al., 1989). Relative to encoding under full attention conditions, dividing attention at encoding impairs free recall, cued recall, and recognition memory performance (Anderson et al., 1998; Baddeley et al., 1984; Craik et al.,1996, 2018; Fernandes & Moscovitch, 2000; Naveh-Benjamin & Guez, 2000). Moreover, memory performance costs under divided attention are greater for contextual memory details compared to item memory, especially when spatial location is used as the context memory task (Troyer et al., 1999; Troyer & Craik, 2000). In contrast, dividing attention at retrieval has minimal effects on memory performance (Craik et al., 1996; Naveh-Benjamin et al., 1998; Naveh-Benjamin et al., 2006). These results have been interpreted to suggest that memory encoding operations are especially demanding of attentional resources.

The exact mechanisms by which attention shapes episodic memory encoding have yet to be fully uncovered. However, emerging evidence from the neuroimaging literature demonstrates that the brain state occurring prior to a presentation of a stimulus, impacts memory formation for that stimulus (Fernandez et al., 1999; Guderian et al., 2009; Otten et al., 2006; Turk-Browne et al., 2006; Yoo et al., 2012). For example, functional connectivity between parietal brain regions mediating attention, and posterior perceptual cortices at encoding was stronger during cue presentations associated with remembered versus forgotten items (Uncapher et al., 2011). This suggests that attention may dictate whether a stimulus would be successfully encoded during a preparatory period even before the stimulus is presented. Yet, fewer studies have investigated the behavioural markers of pre-stimulus attention in relation to episodic memory formation.

Researchers have typically used variations of the sustained attention to response task (SART; Robertson et al., 1997) to investigate how pre-stimulus attentional state affects memory encoding. Traditional SART requires participants to press a button in response to a frequent non-target (GO) stimulus, and withhold button pressing when a non-frequent target (NOGO) stimulus appears on the screen. Failure to suppress responses to NOGO stimuli has been used to index attention failures. A hallmark feature of the SART is an acceleration of RT to the NOGO stimulus preceding an error, followed by a deceleration of RT to the GO stimulus following an error. This variation in RT arguably reflects the disengagement of attention from task-relevant information, and re-direction of attentional resources to the task at hand respectively (Manly et al., 1999; Robertson et al., 1997). One study used a variation of the SART to investigate how fluctuations in attention during encoding impacted verbal memory performance (Smallwood et al., 2006). In this study, participants were shown a sequence of frequent non-target words (GO) for which they were asked to respond to, and non-frequent target words (NOGO) for which they were asked to withhold response. Participants were either assigned to an incidental or intentional encoding condition for the words presented. Retrieval performance was measured via cued recall and results were analysed by a process dissociation method (Jacoby, 1998) to distinguish whether words were retrieved based on recollection or familiarity. Regardless of group assignment, participants showed no difference in probability of recollection- or familiarity-based recall prior to an error (i.e., responding to a NOGO stimulus). However, they were more likely to retrieve words based on recollection after an error has been made, presumably reflecting re-direction of attention at encoding to task-relevant information, thereby increasing chances of recollection-based recall (Smallwood et al., 2006). Another study examined the effect of spontaneous fluctuations in levels of attention on incidental encoding performance (deBettencourt et al., 2018). Participants were shown a series of images sequentially without delay, each for 1 sec. For each image, they were asked to make a categorical decision as fast as possible via button pressing to indicate whether the image reflects an indoor or an outdoor scene. The task was designed so that 90% of the images came from one category (e.g., outdoor), and 10% came from the other category, thereby ensuring that the task was repetitive in nature.

Similar to a typical SART task, participants may get accustomed to responding to the frequent GO stimulus (i.e., outdoor) when attention had presumably been disengaged from the task as reflected by relatively quicker RTs, and therefore may be more likely to commit an error of commission (i.e., pressing outdoor for an indoor stimulus). In contrast, it was predicted that when participants’ attention was directed on the task, they would be less likely to exhibit faster RTs and less likely to commit errors on the task. Indeed, the results confirmed these predictions. More importantly, when retrieval performance for the infrequent NOGO stimuli was assessed the stimuli were more likely to be forgotten if the average RT of the preceding three stimuli was relatively quick, presumably reflecting a pre-stimulus lapse in attention. Based on these results, the authors created a real-time adaptive task to systematically present stimuli from the infrequent category when the attention state of the participant may have dipped below a certain threshold. This threshold was based on a running RT average of the three stimuli preceding a certain stimulus. Results were replicated, and memory for the non-frequent stimuli was worse when participants’ pre-stimulus attention was in a compromised state (deBettencourt et al., 2018).

Results from studies discussed above that combined SART with traditional memory paradigms, support the idea that the ebb and flow of attentional state can influence item memory accuracy even before the items are presented. Yet, it remains unclear how pre-stimulus attention fluctuations at encoding may impact context memory performance. Since encoding contextual details related to an item is more effortful and requires greater attentional resources compared to item memory recognition (Troyer et al., 1999; Troyer & Craik, 2000), it is likely that spontaneous dips in attention levels prior to stimulus presentation may differentially impact context memory relative to item memory, however this assumption has not yet been tested to our knowledge. Additionally, it is important to note that results from the above studies combining adaptive SART and memory paradigms must be interpreted with caution. Criticism has been raised over the potential influence of speed-accuracy-trade-offs regarding the SART task. That is, errors of commission on the SART (responding to a NOGO stimulus), may in fact reflect individual differences in response speed strategies rather than compromised attentional state per se (Helton, 2009; Helton et al., 2009; Seli et al., 2012; Seli, et al., 2013a; Seli, et al., 2013b). For example, Seli et al., (2013b) demonstrated that errors of commission on the SART varied systematically with manipulated differences in response delay, in that, slower responses generated fewer errors. This suggests that SART errors may not merely reflect failures in attention per se, but are significantly impacted by individual differences in speed-accuracy-trade-offs. Finally, while SART allows for investigating how pre-stimulus attention levels impacts encoding success, the task only provides a behavioural marker for pre-stimulus attention on non-frequent target events rather than on every event, thus, it may not capture pre-stimulus attentional fluctuations on a trial-by-trial basis.

In addition to fluctuations in pre-stimulus attentional state, successful memory formation during encoding may also be impacted by attentional shifts away from task-relevant information immediately after encoding (i.e., post-stimulus attentional state). This may interfere with the ongoing consolidation of recently learned information, thereby hampering memory recall (Craig et al., 2014; Dewar et al., 2012). For example, Craig et al. (2014) presented participants with a wordlist followed by a 9-minute rest period where they were cued to think of personal autobiographical memories. Their memory performance for the word list was significantly worse as compared to when the encoding phase was followed by a brief period of quiet wakeful rest. Therefore, it is reasonable to predict that post-encoding consolidation processes may be impacted by spontaneous shifts in attention. However, to our knowledge, no studies have used an explicit online measure to assess the impact of variations of attention on post-encoding consolidation on a trial-by-trial basis.

In the current study, we created a novel task which we refer to as the *Montreal Attention at Encoding Task* (MAET). This task was based off the commonly used psychomotor vigilance task (PVT: Dinges & Powell, 1985), which assesses vigilance/sustained attention and has been used extensively in sleep deprivation studies. We adapted the PVT in the current study to investigate whether and how spontaneous fluctuations in pre- and post-stimulus attention levels during memory encoding, impact memory performance for items and their associative spatial context on a trial-by-trial basis. A typical PVT task measures fluctuation in attention by recording RTs to visual or auditory stimuli that occur frequently at random intervals over a period of time. Relatively longer RTs on a trial-by-trial basis have been suggested to reflect instances where the participant’s attention may have lapsed (Dorrian et al., 2004; Jung et al., 2011; Lim & Dinges, 2008). In the current MAET task, we ask participants to encode a series of object images presented either on the left or right side of the screen. Staggered between the object stimulus presentations is a fixation cross that increases in size after a variable duration. Participants are instructed to respond as quickly as possible to the change in fixation cross size. We operationalize the state of pre-stimulus attention as RTs to the fixation cross preceding an object stimulus. By the same token, the state of post-stimulus attention can be inferred from RTs to the fixation cross following that object stimulus. We use RTs indexing pre- and post-attentional state to predict subsequent memory performance for the objects and their spatial context. We predict that relatively longer pre-stimulus RTs indexing spontaneous dips in attention, will incur greater costs in memory failures for spatial context memory compared to item memory. If spontaneous drifts in attention do impact post-encoding consolidation operations, then we predict that relatively longer post-stimulus RTs indexing momentary reductions in attention will be associated with worse memory performance especially for spatial contextual details related to an item.

## 2. Methods

### 2.1 Participants

Thirty-four healthy young adults (21-34 yrs) successfully completed this study. A statistical power analysis was performed for sample size estimation, based on previous investigations of behavioural effects of attention and memory performance at encoding (Troyer & Craik, 2000) (N = 24). The effect size (cohen’d valaue) was 1.49, considered to be very large using Cohen’s criteria (1988). With an alpha = .05 and power = .08, the projected sample size needed with this effect size is approximately N = 12 for this within-group comparison. Thus, our sample size of 34 will be more than adequate for the main objectives of this study and should allow some buffer for expected post-hoc exclusion. A total of four participants were excluded from our analysis post-hoc. Three participants were excluded due to missing too many responses at encoding, and one participant was excluded for having an unusually slow mean response time (RT) at encoding. Post-hoc exclusion was based on meeting the 1.5 * Inter quartile range outlier rule. The final sample consisted of 30 participants (age range 21-34 yrs, mean age = 25.76 yrs; 17 females; mean years of formal education [EDU] = 15.80 yrs). The MAET was adapted for functional magnetic resonance (fMRI) testing to examine the neural underpinnings of pre- and post-stimulus attention and episodic memory using this novel paradigm. All eligible participants underwent fMRI brain scanning while performing both encoding and retrieval phases of the task. The behavioural results of the task will be discussed in this manuscript while the neuroimaging results will be discussed in future publications.

The study involved two sessions, each conducted on a separate day. Session 1 consisted of a medical questionnaire, a battery of neuropsychological tests and a practice run of the MAET performed in a mock MRI scanner. The Beck Depression Inventory (BDI-II), exclusion cut-off > 14, (Beck et al., 1996) and the California Verbal Learning Test (CVLT) Delay Free Recall (DFR), exclusion cut-off < 11 (Mervielde et al., 1999) were used to assess eligibility. Additional exclusion criteria based on the medical questionnaire responses included: having a history of psychiatric illness, substance abuse, and/or neurological insult resulting in a loss of consciousness of > 1 minute. During session 1, participants also completed the CVLT Delay Cued Recall (DCR), and CVLT Delay Recognition (DRG) to assess long term verbal memory (Mervielde et al., 1999). To assess executive function, the Wisconsin Card Sorting Test: computer version 4 (WCST; Heaton & PAR Staff, 2003) and the Delis-Kaplan Executive Function System (D-KEFS; Delis, Kaplan, & Kramer, 2001) verbal fluency test; Letter Fluency (LF), Category Fluency (CF), and Category Switching (CS), were also administered. To assess the frequency of everyday attentional lapses, the Cognitive Failure Questionnaire (CFQ; Broadbent et al., 1982) and the Mindful Attention Awareness Scale-Lapses Only (MAAS-LO; Carriere et al., 2008) were administered.

Those who met the medical inclusion criteria, BDI-II and the CVLT cut-offs, and performed above chance on the mock-MRI scanner trials, were invited to participate in a second fMRI testing session. All participants were recruited via online and printed advertisements from within the Greater Montreal region. Participants were paid and provided their informed consent to participate in the study. The ethics board of the Douglas Research Centre, Le Centre intégré universitaire de santé et de services sociaux (CIUSSS) de l‘Ouest-de-l’Île-de-Montréal approved the study protocol.

### 2.2 The Montreal Attention at Encoding Task

To explore the effects of pre- and post-stimulus attention on item and spatial context memory performance, we created the MAET task (see figure 1). Participants were asked to encode pictures of objects and their location on each trial, and to respond to a fixation cross that expanded in size after a random duration. They were then subsequently presented with a retrieval run consisting of an equal number of old and new objects intermixed at random, and their memory for the objects previously seen was tested. In total, participants completed 4 experimental runs of encoding and 4 runs of retrieval. Each encoding run was subsequently followed by its respective retrieval run with a brief distraction task between the encoding and retrieval phases (details discussed below). The order of run presentation was counterbalanced across participants.

**Figure 1.**
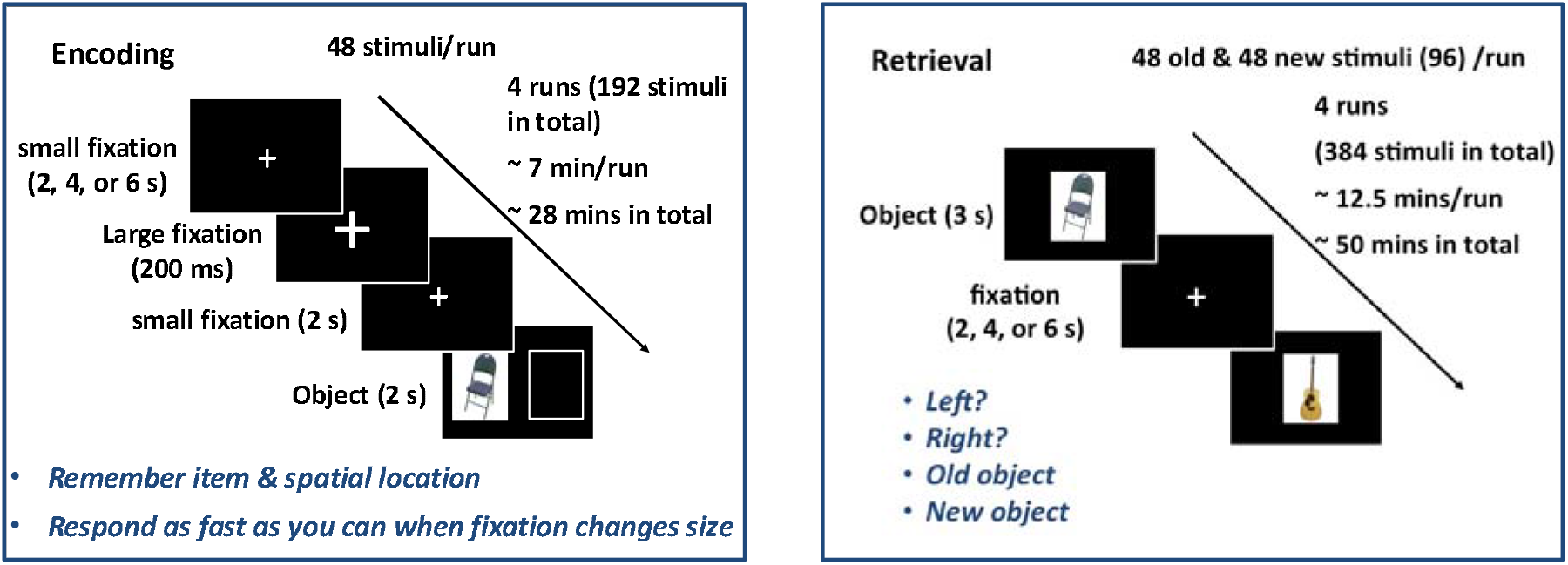
MAET task procedure. MAET included an encoding and retrieval phase. On each encoding trial, participants were presented with a fixation cross that expanded in size and were asked to respond to the change in size as quickly as possible. An object stimulus to be encoded followed either on the left or right side of the screen and participants had to encode the object and its location. Response times to the fixation cross were used to attention levels on a trial-by-trial basis. On each retrieval trial, a centrally presented old or new object was shown and participants had to indicate whether the object was previously presented on the left, right, whether it was an old object but they don’t remember its initial location, or whether it’s a new object.

The task stimuli set was obtained from the freely available Bank of Standardized Stimuli (BOSS; Brodeur et al., 2014), which offers high quality and standardized images of objects, and has been used in several other behavioural and neuroimaging studies. Each encoding run consisted of objects (48 in total) randomly presented either on the left (24 objects) or right (24 objects) side of the screen. These objects were equally selected from 12 different categories (i.e.; food, musical instruments, vehicles, building infrastructure, clothing, decoration and gift, electronics, game and entertainment, household articles, kitchen items, outdoor and sport equipment, and stationary items). Similarly, each retrieval run consisted of a balanced distribution of objects selected from the same 12 categories (48 old and 48 new objects). Upon completion of all runs of the task, participants were asked to complete the Thinking Content component of the Dundee Stress State Questionnaire (DSSQ; Mervielde et al., 1999) to assess mind-wandering during the task. The task was programmed and run using E-Prime (Psychology Software Tools, Inc.; Pittsburgh, PA, USA) software. Participants requiring visual acuity correction wore corrective plastic lenses and two fiber optic 4-button response boxes were supplied to participants to make responses during the task. To ensure that responding to the fixation cross was indeed a valid index of attention, we computed an error rate for each participant (i.e., trials where participants failed to respond to the fixation cross divided by total number of encoding trials completed) and correlated the error rate with accuracy on the task, and the attentional lapses questionnaires (CFQ, DSSQ, and MASS-LO).

### 2.2.1 Encoding phase

Each encoding trial commenced with a small fixation cross which was presented in the middle of the screen and increased in size after a random duration (2, 4, or 6 secs). The large fixation cross lasted on the screen for 200 msec then collapsed back to its original size and lasted for another 2 secs before commencing a new trial. Participants were asked to press a button as fast as possible when they detect the change in fixation cross size. This was followed by an object randomly presented either on the left or right side of the screen for 2 secs. Participants had to memorize the object and its location. Therefore, the instructions were to memorize the objects and their location, and to respond to the change in fixation cross size. These two sets of instructions were emphasized equally prior to beginning each run. The initial small variable fixation cross served to introduce jitter to the fMRI acquisition, and to prevent stimulus expectancy effects similar to the PVT (Dinges & Powell, 1985). The variability in RTs to the fixation cross size change were used to gauge attention levels. The encoding phase consisted of 4 runs (48 stimuli/run) for a total of 192 stimuli across all runs. Following each encoding run, participants completed a short distraction task (60 secs) to minimize rehearsal of encoded information before commencing the retrieval run. During the distraction phase, participants were simultaneously shown two words and were asked to select the word that comes first in the alphabet.

### 2.2.1 Retrieval phase

On each retrieval trial, participants were shown either an old (previously presented during encoding), or new object presented centrally for 3 secs. For each object they were asked to indicate whether the object was either i) previously presented on the left side; ii) previously presented on the right side; iii) an old object, but not sure of its initial location; iv) a new object. Trials were separated by a variable ITI fixation (2, 4, or 6 secs). This design allowed for the distinction between associative context retrieval and item memory retrieval without its spatial context. Each retrieval run consisted of 96 objects (48 old and 48 new) for a total of 384 objects across all 4 retrieval runs.

### 2.3 Behavioural data analysis

We calculated mean accuracy for all possible response types:

1. Associative context hits: Correctly recalling an object and its spatial location.

2. Item memory: Recognizing old objects but providing no associative spatial location.

3. Context misattribution: Recognizing old objects but providing incorrect associative spatial location (e.g., saying an object previously seen on the left was initially presented on the right).

4. Miss: Incorrectly identifying old objects as new.

5. Correct rejection: Correctly identifying new objects.

6. False alarm: Incorrectly identifying new objects as old.

The percentage of associative context hits, item memory and context misattribution were calculated as the number of responses in each respective category, divided by the total number of old hits completed per participant. The percentage of correct rejection and false alarms were calculated as the number of trials in each of the two categories, divided by the total number of new trials completed per participant. Trials where participants failed to respond at retrieval (3.31% out of the total retrieval trials completed) were excluded from the analysis. We computed overall accuracy (i.e., hits) as the sum of associative context hits, item memory and context misattribution, divided by the total number of old trials completed per participant. We also computed proportion associative context hits (correct associative context trials/total no. of hits), and proportion item memory (item memory trials/total no. of hits). D’ was used as a measure of sensitivity and was computed as overall standardized hit rate minus standardized false alarm rate.

To ensure that participants scored above chance level, we conducted one-sample t-tests on hits, associative hits, and correct rejection. In addition to using d’ as a measure of response discriminability, we calculated chance level based on response probability (i.e., probability of selecting the correct response by chance), and stimulus presentation probability (i.e., probability of a stimulus type appearing by chance) (Snytte et al., 2020). For example, response probability for hits = 75% (three out of 4 button responses would lead to a general hit i.e., old-left, old-right, old with no associated spatial location). On the other hand, stimulus presentation probability for hits would equal 50% (half of the stimuli presented at retrieval were old). Therefore, by compounding response and stimulus probability for hits, chance level would equal 37.5%. Chance level for correctly identifying an old item previously shown on the left or right was 6.25%. To calculate the overall chance level for associative context hits, we computed the sum of both probabilities (6.25% + 6.25%), which yielded 12.5%. Similar to associative context hits, chance level for correct rejection was computed as 12.5%.

### 2.3.1 Logistic regression analysis

We used logistic regression analysis to test whether pre- and post-stimulus attention levels differentially impact associative context versus item memory accuracy within-subject. Pre-stimulus attention levels on a trial-by-trial basis were indexed via RTs to the fixation cross preceding each object stimulus to be encoded, while post-stimulus attention levels were indexed via RTs to the fixation cross following each object. For subsequent memory analysis, correct associative context trials were scored as 1, and item memory was scored as 0. Therefore, each encoding trial was associated with its respective pre-stimulus RT, post-stimulus RT and subsequent memory score (i.e., 1 for correct associative context and 0 for item memory). Trials with missing responses at encoding or retrieval, context misattribution trials, and miss trials were excluded from analysis. For each subject, a logistic regression analysis was conducted to predict the binary memory variable (i.e., 1, 0) from the RT index of pre-stimulus attention level during encoding. Another logistic regression analysis was conducted to predict the same binary memory variable from the RT index of post-stimulus attention level during encoding for each subject. Finally, we compared the resulting beta values for all participants via one sample t-test to determine their overall direction at the group level.

We were also interested in testing whether fluctuations in pre- and post-stimulus attention levels at encoding impacts associative context recollection more broadly. To that end, we ran a third within-participant logistic regression model testing whether pre-stimulus attention levels as indexed by pre-stimulus fixation RTs, would predict associative context hits, versus associative context failure (i.e., collapsing item memory, context misattribution, and miss events). The resulting beta values for all participants were then compared via one sample t-test to determine their overall direction at the group level. Lastly, we ran a fourth and final logistic regression model to predict associative context hits, versus associative context failure (i.e., collapsing item memory, context misattribution, and miss events) within-participant, from post stimulus RTs at encoding. The resulting beta values were then compared via one sample t-tests to determine their overall direction at the group level.

## 3. Results

### 3.1 Accuracy and reaction time results

Table 1. summarizes demographics and neuropsychological test data, and figure 2a shows a violin plot of accuracy results on the MAET. Participants scored well above chance level on associative context hits (M = .56, SD = .20, t(29) = 12.17, p < .001, hits (M = .78, SD = .14, t(29) = 15.77, p < .001), and correct rejection (M = .85, SD = .20, t(29) = 34.57, p < .001). Means and SDs of accuracy results are presented in Table 2.

**Table 1.** Demographics and neuropsychological test data

|  |  |
| --- | --- |
| **Sample size** | 30 |
| **Age (Yrs)** | 25.76 (0.77) |
| **Gender (n, [%] females)** | 17 [57%] |
| **EDU (Yrs)** | 15.80 (0.35) |
| **CVLT – DFR<sup>b</sup>** | 12.63 (0.32) |
| **CVLT – DCR** | 12.87 (0.36) |
| **CVLT – DRG** | 15.23 (0.18) |
| **WCST – categories completed<sup>c</sup>** | 8.00 (0.27) |
| **WCST – % correct<sup>c</sup>** | 0.81 (1.88) |
| **D-KEFS – LF** | 37.20 (1.71) |
| **D-KEFS – CF<sup>d</sup>** | 35.83 (1.63) |
| **D-KEFS – CS** | 14.23 (0.51) |
| **Estimated IQ (AMNART)** | 112.80 (1.45) |
| **CFQ – Total score** | 29.63 (1.99) |
| **CFQ – Forgetfulness** | 10.57 (0.82) |
| **CFQ – Distractibility** | 11.10 (0.86) |
| **CFQ – False triggering** | 6.67 (0.62) |
| **MASS – Lapses** | 30.33 (1.33) |
| **DSSQ – TRI** | 20.53 (1.08) |
| **DSSQ – TUI** | 14.63 (0.94) |

Note: This table presents means and standard deviations between brackets for demographic, neuropsychological measures. EDU = Years of Education; CVLT = California Verbal Learning Test; DFR = Delay Free Recall; DCR = Delay Cued Recall; DRG = Delay Recognition; WCST = Wisconsin Card Sorting test; D-KEFS = Delis-Kaplan Executive Function System; LF = Letter Fluency; CF = Category Fluency; CS = Category switching; AMNART = American National Adult Reading Test; CFQ = Cognitive failures Questionnaire; MASS = Mindful Attention Awareness Scale (lapses only); DSSQ = Dundee Stress State Questionnaire; TRI = Task-related Interference; TUI = Task-unrelated Interference.

**Table 2.** MAET task accuracy results

|  |  |
| --- | --- |
| **Hits** | .78 (.14) |
| **Associative context hits** | .56 (.20) |
| **Proportion associative context hits** | .68 (.16) |
| **Item memory** | .19 (.10) |
| **Proportion Item memory** | .25 (.15) |
| **Context misattribution** | .05 (.04) |
| **Correct rejection** | .85 (.20) |
| **False alarms** | .15 (.12) |
| **Misses** | .20 (.14) |
| **d'** | 2.08 (.97) |

Note: This table presents means and standard deviations between brackets for accuracy results for the MAET task.

**Figure 2.**
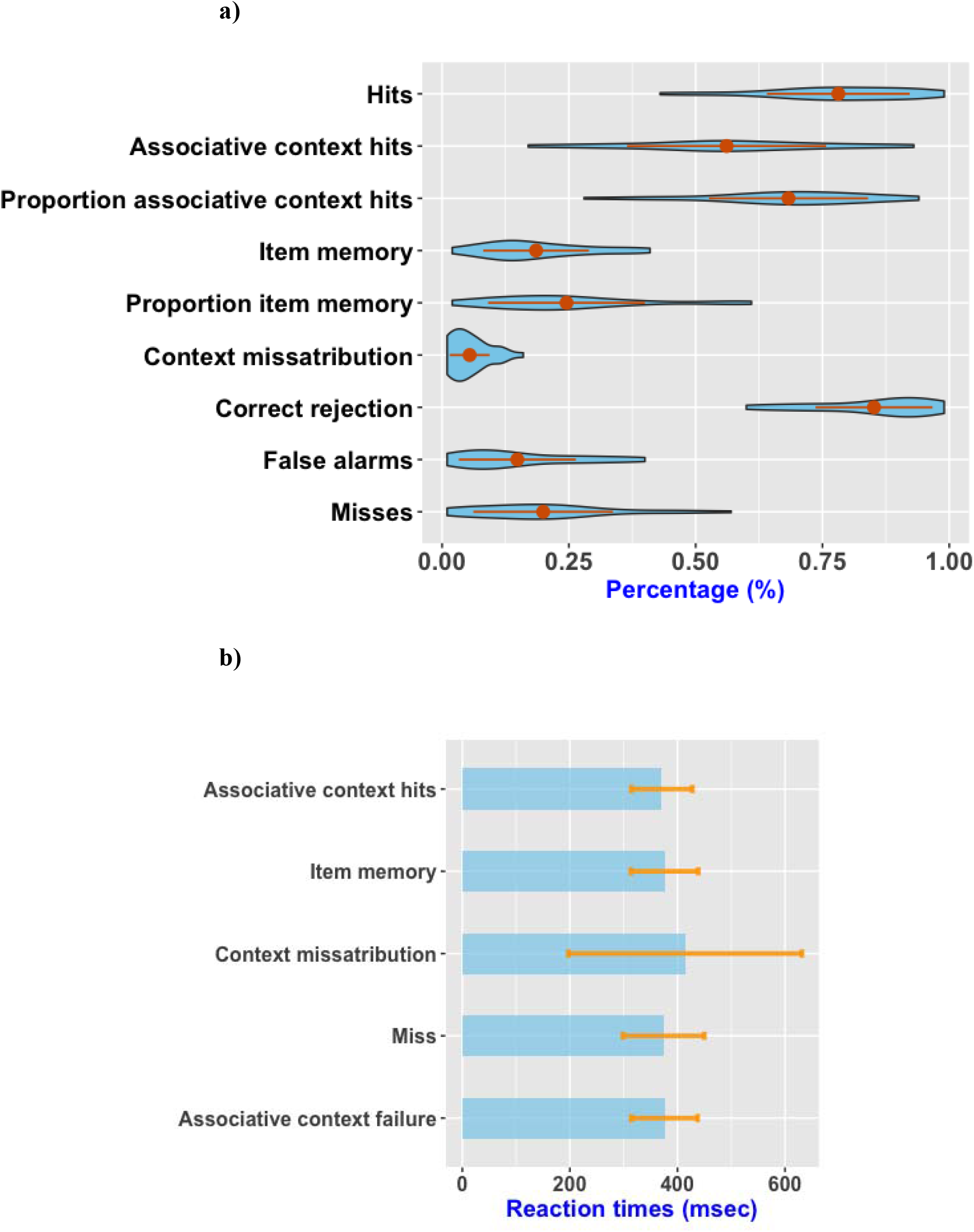
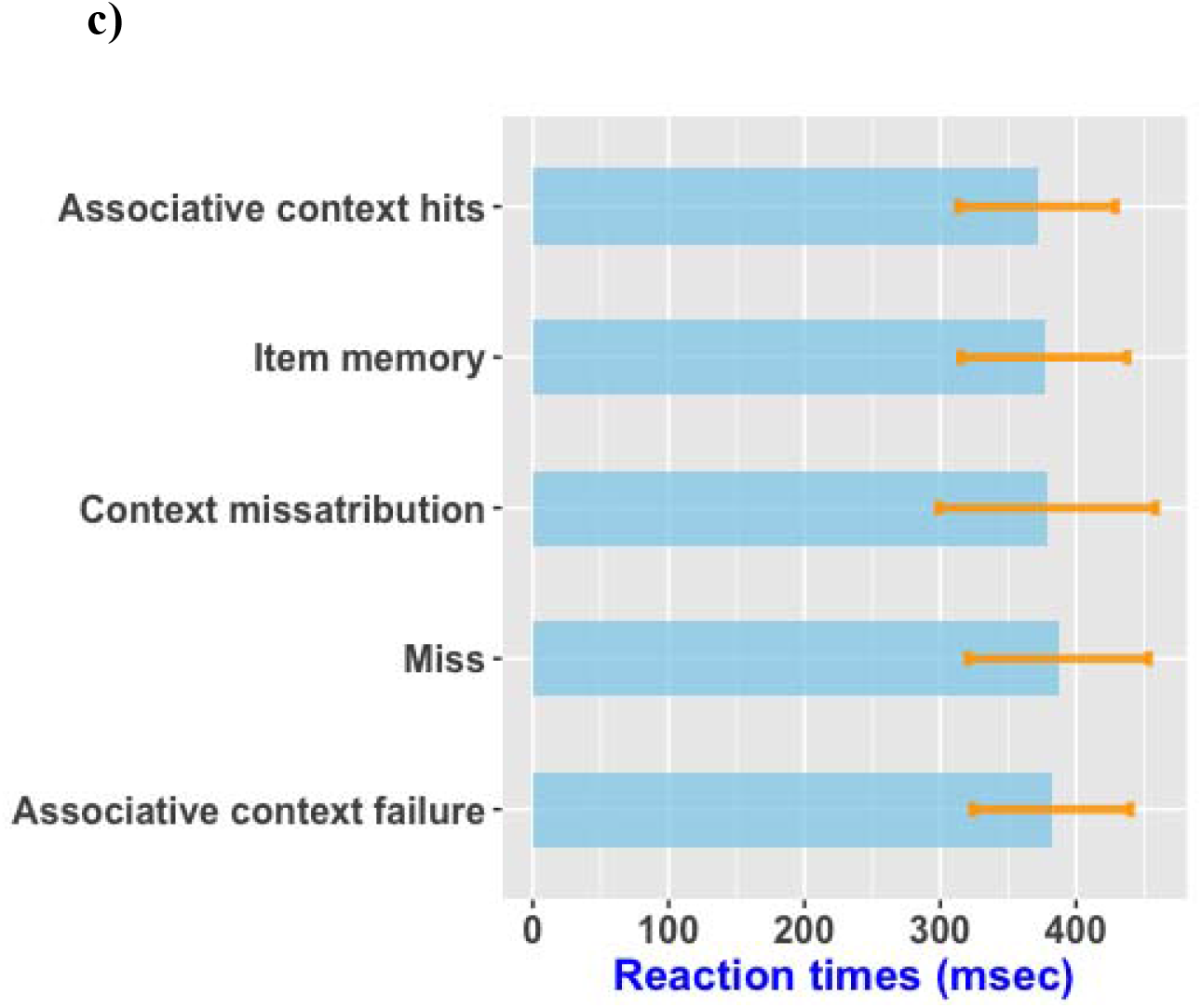
Plots of accuracy and response times for MAET task. a) Violin plot showing mean accuracy results and standard deviations for each event category on the MAET task. Participants scored well above chance level on hits, associative context hits, and correct rejection. Associative context memory hits percentage was significantly greater than item memory. b) Bar plot showing mean response time and standard deviations to the variable fixation cross presented prior to stimuli that were subsequently scored as associative context hit, item memory, context misattribution, miss, and associative context memory failure (i.e, collapsing item memory, context misattribution, and miss events). The mean response times for the different stimulus categories were not significantly different from each other. c) Bar plot showing mean response time and standard deviations to the variable fixation cross presented after stimuli that were subsequently scored as associative context hit, item memory, context misattribution, miss, and associative context memory failure (i.e, collapsing item memory, context misattribution, and miss events). The mean response times for the different stimulus categories were not significantly different from each other.

The mean RT to the fixation cross staggered between the object stimuli at encoding was 373.68 msec (SD = 57.51). Figure 2b displays a bar plot of mean pre-stimulus RT to objects that were subsequently scored as associative context hit, item memory, context misattribution, miss, and associative context failure (i.e., collapsing item memory, context misattribution, and miss events). Figure 2c displays a bar plot of mean post-stimulus RT to objects that were subsequently scored as associative context hit, item memory, context misattribution, miss, and associative context failure. Post-hoc tests revealed that mean pre-stimulus RTs associated with the different stimulus categories were not significantly different from each other. Similarly, mean post-stimulus RTs associated with the different stimulus categories were not significantly different from each other. There were no sex differences across participants in MAET memory performance or mean RT of fixation cross at encoding (ps > .05). Overall, participants performed the task well as indicated by a relatively low error rate (M = 2.34%, SD = 2.80) and high d’ value (M = 2.08, SD = 0.97) reflecting high task sensitivity. Error rate was not associated with task accuracy, yet, it was positively correlated with the distractibility subscale on the CFQ (p = .038) and showed a marginally significant positive correlation with task unrelated interference – DSSQ (p = .078).

### 3.2 Regression analysis results

We first evaluated the relationship between associative context and item memory accuracy at retrieval, and our RT index of pre-stimulus attentional state at encoding. We used logistic regression to evaluate this relationship within-participant, and then compared the resulting beta values to determine their overall direction at the group level. Shapiro-Wilks test showed that the data did not violate assumptions of normality (p = .92), and one sample t-test performed at the group level revealed that the overall direction of the betas was not different than zero (p = .72). Therefore, this regression analysis failed to support our hypothesis that pre-stimulus fluctuations in levels of attention at encoding predict associative context and item memory at retrieval. We ran another logistic regression model testing whether fluctuations in pre-stimulus attention levels at encoding would predict associative context memory more broadly (i.e., associative context success vs. associative context failure). Shapiro-Wilks test showed that the data did not violate assumptions of normality (p = .84), and one sample t-test performed at the group level revealed that the overall direction of the betas was not different than zero (p = .09). These results suggest that fluctuations in pre-stimulus attention levels as indexed by RTs to fixations presented prior to object stimuli to be encoded, did not impact associative context vs. item memory performance, nor did they predict associative context hits versus associative context failure more broadly.

Similar to the analyses outlined above, we ran within-participant logistic regressions to evaluate whether post-stimulus RTs predicted associative context hits vs. item memory, and then compared the resulting beta values to determine their overall direction at the group level. Shapiro-Wilks test showed that the data did not violate assumptions of normality (p = .07), and one sample t-test performed at the group level revealed that the overall direction of the betas was not different than zero (p = .18). Hence, post-stimulus RTs failed to predict associative context hits vs. item memory. Our final within-participant logistic regression analysis tested whether post-stimulus RTs would predict associative context hits vs. associative context failure more broadly. Our results revealed that the majority of the participants (20/30) showed negative beta values, reflecting that quicker post-stimulus RTs at encoding predicted associative context hits, and slower post-stimulus RTs predicted associative context failure. To test whether this relationship holds at the group level, we ran a one sample t-test to determine the overall direction of the beta values after making sure that the data did not violate assumptions of normality (Shapiro-Wilks test: p = .43). The logistic functions for all participants are plotted in figure 3, revealing a reliably negative slope on average (p = .01), indicating that faster RT to the fixation cross that ensued the object stimuli was positively correlated with associative context memory success, and slower post-stimulus RTs predicted associative context failure at retrieval. We also compared the two groups of participants showing positive (n=10), and negative (n=20) slopes on MAET memory performance. Results showed that participants with negative slopes had higher hits rate (t(28) = −2.59, p = .02), and associative context memory accuracy (t(28) = −2.20, p = .03), compared to those with positive slopes.

**Figure 3.**
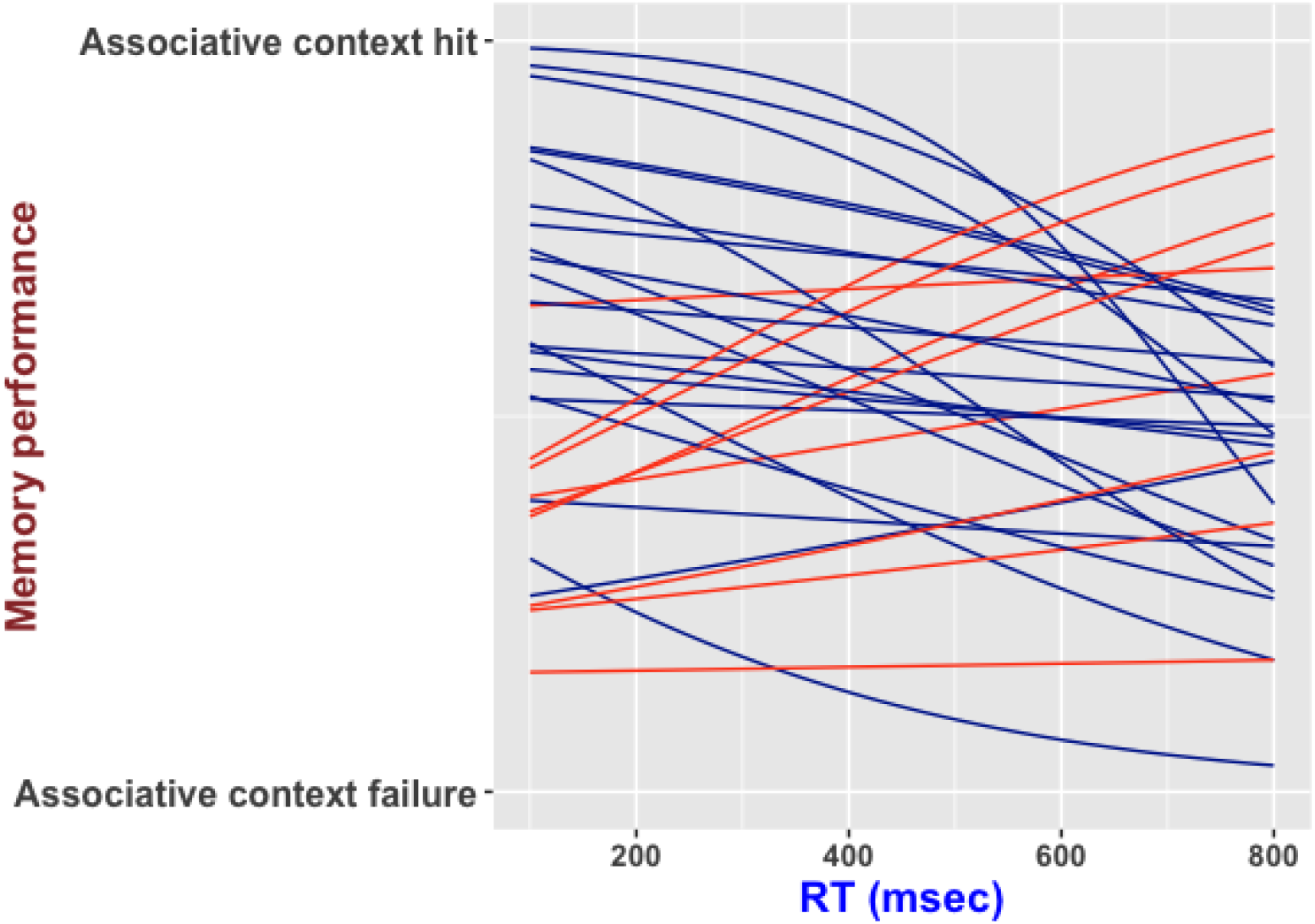
Logistic functions of all participants predicting associative context success vs. associative context failure from post-stimulus RTs. The logistic function for each participant is plotted on the same graph to visualize the relationship between post-stimulus encoding response times, and associative context success vs. associative context failure. Red lines demonstrate logistic functions with positive slopes and blue lines demonstrate logistic functions with negative slopes. As indicated on the graph, the majority of participants (20/30) had negative slopes indicating a reliably negative slope on average (p = .025). That is, faster post-stimulus response times at encoding predicted associative context memory success, and slower post-stimulus response times predicted associative context memory failure.

### 4. Discussion

In the current study, we designed a novel task (MAET) to examine how moment-by-moment fluctuations in pre- and post-stimulus attentional state at memory encoding, may impact memory performance for items and their contextual details. Pre-stimulus attention level at encoding was gauged via RT to a fixation cross presented prior to the object stimulus to be encoded on a trial-by-trial basis. Conversely, post-stimulus attention level on a trial-by-trial basis was indexed via RT to a fixation cross following each object stimulus at encoding. Our results revealed that pre-stimulus attentional state at encoding as indicated by RTs, did not predict associative context memory vs. item memory, nor did it predict context memory success more broadly. On the other hand, context memory encoding success was associated with post-stimulus RTs. We discuss these results and their implications in more detail below.

Contrary to dual-task, divided attention paradigms that require participants to concurrently and simultaneously encode items into memory while engaging in a secondary task, the MAET employed in the current study required participants to sequentially respond to a variable fixation cross and encode objects into memory. Hence, this task can be considered a task-switching paradigm whereby two sets of stimuli are presented sequentially without temporal overlap (Kiesel et al., 2010; Monsell, 2003). We recently employed the same task in our laboratory in a different study without the task-switching component (i.e., participants only had to memorize the stimuli without having to respond to a variable fixation) in a sample of healthy young adults (Snytte et al., 2020), and memory performance was comparable to that of the current study. Therefore, despite the task-switching nature of the current paradigm, task-switching costs were minimal, and likely did not affect memory performance. Moreover, unlike SART, the MAET task is not purely a task of vigilance/sustained attention. Switching between the encoding stimulus and responding to the fixation cross likely disrupts the vigil. However, error rates in the current task predicted lapses in attention as measured via the CFQ scale (Broadbent et al., 1982), and showed a marginal positive association with task unrelated interference or mind-wandering (Mervielde et al., 1999). Hence, our task is sensitive to the experience of everyday attentional lapses, and may also be sensitive to mind-wandering similar to SART (Robertson et al., 1997; Smallwood et al., 2004).

Previous studies have demonstrated that encoding attentional state prior to episodic events impacts subsequent memory for those events (deBettencourt et al., 2018; Markant et al., 2014). Moreover studies using divided attention paradigms have demonstrated that when attention is divided during encoding, memory for contextual details is differentially impaired relative to item memory, presumably due to the greater attentional demands required for context memory encoding (Troyer et al., 1999; Troyer & Craik, 2000). As such, we predicted that spontaneous fluctuations in levels of pre-stimulus attention at encoding, would differentially impact context memory performance relative to item memory. Surprisingly, results from our logistic regression analysis did not support our hypothesis. In a secondary logistic regression analysis, we further tested whether fluctuations in pre-stimulus attentional state would predict associative context success vs. failure more broadly, however this analysis also failed to reveal significant effects. Taken together, fluctuations in pre-stimulus attentional state as indexed by RTs to a fixation cross presented prior to the object stimulus to be encoded in the MAET task, did not impact context memory encoding success. The lack of association between pre-stimulus RTs and associative context vs. item memory may indeed reflect the insensitivity of context memory encoding to fluctuations in pre-stimulus attention state. Yet, another possible explanation points to the low statistical power in this specific analysis. Our sample consisted of healthy young adults who performed the task generally well, scoring well above chance level and selecting the correct associative context on more than half of the trials. However, by virtue of our task design, more correct associative context trials would result in fewer item memory trials, hence creating an unbalanced number of events between the two stimulus categories. Indeed, there were significantly more correct associative context trials relative to item memory trials, which may have reduced the power in our logistic regression analysis. It is also possible that participants were aware of their performance on the task, such that relatively longer pre-stimulus RTs may have signalled that their attention was drifting away from the task. Consequently, this may have triggered the re-direction of attentional resources to task-relevant information, offsetting the effects of momentary dips in pre-stimulus attention, thereby not compromising context memory encoding performance. While our results may seem to suggest that pre-stimulus attention levels do not impact context memory encoding performance when taken at face value, these findings should be taken with a grain of salt given our novel task design and the task limitations discussed above.

Given the versatility of our MAET task design, we also investigated whether RTs presented immediately after an object encoding event would predict associative context vs. item memory or associative context hits vs. associative context failure in general. The variable fixation cross staggered between the object stimuli at encoding lasted 4 secs on average before expanding in size and a behavioural response was made. Therefore, RT to a fixation cross presented after an object stimulus potentially reflects the attentional state of a participant during that object presentation or moments after (i.e., post-stimulus attention) leading to the subsequent behavioural response. Our results revealed that post-stimulus attention did not predict context vs. item memory but did in fact predict associative context memory encoding success more generally. That is, relatively longer post-stimulus RTs, presumably signalling brief interruptions in attention either during the encoding event itself or moments after predicted context memory failure, and relatively shorter post-stimulus attention predicted context memory hits.

Overall, our findings suggest that there are attention states where individuals are optimally tuned in for successful context memory encoding and other states where contextual details are likely to be missed. A possible mediator of these effects is mind-wandering, or the shift of attention from task-related processes to self-generated processes unrelated to the task (Smallwood & Schooler, 2006). In the current study, mind wandering, was only marginally related to error rates (p = .078) on the task and was not directly linked to memory performance. However, mind-wandering in the current study was assessed via a self-report questionnaire presented at the end of the study, and therefore may not have captured the moment-by-moment fluctuations in attention that occurred during the encoding task. Furthermore, evidence linking mind-wandering to task performance is inconsistent throughout the literature (McVay & Kane, 2012; Seli et al., 2015). Task-unrelated thoughts (i.e., mind-wandering) could also be assessed via thought probes presented on a trial-by-trial basis (Seli, Risko, & Smilek, 2016; Seli et al., 2018), however, we opted against this method since it would add another level of task-switching to our paradigm and would likely disrupt the vigil further, making it difficult to conclude whether mind-wandering or task-switching demands mediated task performance. According to mind-wandering theory, task-unrelated thoughts stem from underload, or underarousal associated with a task (Robertson et al., 1997; Seli, Risko, Smilek, et al., 2016; Thomson et al., 2014). Given the high memory performance level on the current task and its low level of complexity, we speculate that our task afforded most participants to disengage their attention from context memory encoding to task unrelated thoughts (i.e., mind-wandering) after viewing each object, hence explaining the positive relationship between slower post-stimulus RTs at encoding and context memory failure. Our results do not directly speak to the exact post-encoding mechanisms that are disrupted in the face of mind-wandering afforded by momentary dips in post-stimulus attention, however, we hypothesize that disruption likely occurs during the automatic consolidation of recently acquired memory traces that converts those memories into long-term storage (Squire et al., 2015). Disruption of memory consolidation processes could explain the link between momentary fluctuations in attention, mind-wandering, and episodic memory performance in the MAET task, and is in line with previous accounts outlining some of the factors that hinders consolidation of recently acquired information (Craig et al., 2014; Dewar et al., 2012; Mednick et al., 2011). While this hypothesis is highly speculative, future studies utilizing the MAET may help delineate the exact post-encoding mechanisms that are disrupted during momentary ebbs in attention at encoding.

Although most participants in the current study demonstrated a positive relationship between slower post-stimulus RTs at encoding and context memory failure. A few participants demonstrated the opposite pattern of results (i.e., slower post-stimulus RTs linked to context memory success). While it is difficult to explain these results, those participants could potentially be demonstrating speed-accuracy trade-offs. That is, rather than giving equal weight to responding to the fixation cross and encoding a given stimulus, they could be directing more cognitive resources towards the encoding task by rehearsing the object stimuli during the presentation of the fixation cross, thereby responding relatively slow to the fixation stimuli. However, this interpretation is rather unlikely since participants showing a positive relationship between faster post-stimulus RTs and associative context memory success showed more accurate associative context memory performance compared to those who demonstrated the opposite pattern. Further experimentation is required to clarify those results.

In sum, our study introduces a novel paradigm to investigate the influence of the ebb and flow of attentional state on memory performance and demonstrates that context memory success is related to an efficient post-stimulus attention state. Teasing apart the specific post-stimulus cognitive operations that contribute successfully to context memory encoding is worth investigating in future studies to expand the literature on the interaction between the attention and memory systems. Furthermore, understanding the factors related to individual differences in the relationship between momentary fluctuations in attention and memory performance would shed light on the susceptibility of individuals to memory failures, and may offer a unique window for investigating the optimal conditions for learning and memory. Finally, in efforts to pin down the neural mechanisms that underline the relationship between encoding attentional state and context memory success, we have designed and tested the current task to make it suitable for fMRI. We aim to address these neuroimaging results in future publications.

## Acknowledgements

No part of the study procedures or analysis was preregistered in an institutional registry prior to the research being conducted. The conditions of our ethics approval do not permit sharing of the data supporting this study with any individual outside the author team under any circumstances. However, the scripts used to conduct analysis will be made available on Open Science Framework. This work was supported by Natural Science and Engineering Council of Canada (NSERC) # 8400826 awarded to M. N. Rajah. We thank the participants of our study for their time and support in making this project possible.

